# Molecular Screening of Familial Hypercholesterolemia in the Icelandic Population

**DOI:** 10.1101/425975

**Authors:** Greg Kellogg, Bolli Thorsson, Ying Cai, Robert Wisotzkey, Andrew Pollock, Matthew Akana, Rebecca Fox, Michael Jansen, Elias F. Gudmundsson, Bonny Patel, Chihyu Chang, Malgorzata Jaremko, Valur Emilsson, Vilmundur Gudnason, Oscar Puig

**Author notes:** To whom correspondence should be addressed Oscar Puig, PhD, Phosphorus Diagnostics,data. Analytical sensitivity of the tes t1140 Broadway, Suite 1100, New York, 10001, NY, USA.

## Abstract

Familial hypercholesterolemia (FH) is a monogenic disease characterized by a lifelong exposure to high LDL-C levels that can lead to early onset coronary heart disease (CHD). The main causes of FH identified to date include loss-of-function mutations in *LDLR* or *APOB*, or gain-of-function mutations in *PCSK9*. Early diagnosis and genetic testing of FH suspects is critical for improved prognosis of affected individuals as lipid lowering treatments are effective in preventing CHD related morbidity and mortality. In the present manuscript, we developed a comprehensive next generation sequencing (NGS) panel which we applied on two different resources of FH in the Icelandic population: 62 subjects from 23 FH families with known or unknown culprit mutations, and a population-based sampling of 315 subjects selected for total cholesterol levels above the 95^th^ percentile cut-point. The application of the NGS panel revealed significant diagnostic yields in identifying pathogenic *LDLR* mutations in both family and population-based genetic testing.

## Introduction

Familial Hypercholesterolemia (FH, OMIM #143890) is caused by genetic defects in the low-density lipoprotein receptor (*LDLR*) pathway, resulting in lifelong elevation of plasma LDL-cholesterol (LDL-C) concentrations in affected individuals that are largely resistant to caloric restriction, weight loss and physical exercise interventions. At the molecular level, FH can be caused by loss-of-function mutations in *LDLR*, mutations in *APOB* that affect the LDL receptor-binding domain of apolipoprotein B, or gain-of-function mutations in *PCSK9* (1). Mutations in *LDLRAP1* cause Autosomal Recessive Hypercholesterolemia (ARH, OMIM #603813), which has a recessive mode of inheritance and similar clinical manifestation as FH. In summary, FH is a monogenic disease characterized by a lifelong exposure to high LDL-C levels often resulting in premature atherosclerotic cardiovascular disease (2).

A diagnosis of FH can be made on the basis of the plasma total and low density lipoprotein cholesterol concentrations combined with either a clinical examination and family history, or a genetic test. Identification of subjects before the onset of overt cardiovascular disease provides an opportunity to initiate preventive statin therapy (3, 4). However, FH is grossly underdiagnosed and undertreated worldwide (2), and it is estimated that in the US alone one million people remain undiagnosed (5, 6). Early diagnosis and treatment of FH with lipid-lowering therapy has proven to be both cost efficient and effective in mitigating cardiovascular morbidity and mortality risk (7–11).

Traditionally, genetic diagnosis of FH is performed by microarray, PCR or DNA sequencing using Sanger sequencing of all coding regions in *LDLR* plus one or two specific exons in *APOB* (12). Large rearrangements in *LDLR* can be detected by multiplex ligation-dependent probe amplification (MLPA) (13). However, Next Generation Sequencing (NGS) allows parallel sequencing quickly and inexpensively, with high levels of specificity and sensitivity, and several NGS gene panels are already available to diagnose FH (14–17).

In the present study, we performed a molecular screening of FH using two different datasets from the Icelandic population: 1) a cohort of 62 individuals from 23 Icelandic FH families, and 2) 315 individuals with total cholesterol above the 95^th^ percentile cut-point from the Risk Evaluation For INfarct Estimates (REFINE) Reykjavik study (18). REFINE Reykjavik is a prospective cohort study on risk factors and etiology of atherosclerotic disease of 6940 inhabitants from the greater Reykjavik area aged between 20 and 74 years. For the identification of pathogenic mutations causing FH and ARH we used a comprehensive NGS cardiovascular panel targeting the *LDLR*, *APOB*, *PCSK9* and *LDLRAP1* genes.

## Materials and Methods

### Subjects and DNA samples

The Icelandic Heart Association Family Hypercholesterolemia (IHA-FH) study is a family study that aims at identifying all individuals with FH in the Icelandic population. Participants in the IHA-FH study were either family members from previously known FH families identified in the Lipid Research Clinic at the University Hospital of Iceland or individuals admitted to the study from physicians in Iceland (cardiologists and general practitioners) in order to verify clinical diagnosis of FH. Three culprit mutations in the *LDLR* gene have already been identified to date in the Icelandic population (19). A total of 62 members from 23 Icelandic FH families were included in the present study. Here, individuals (N =12 from 3 FH families) that were carriers of any one of these three mutations served as positive controls in this study in order to verify their genetic diagnosis. Further, family members diagnosed free of FH were included as negative controls (N =14, from 11 families). Finally, individuals from the IHA-FH study with clinical diagnosis of FH but unknown culprit mutations were included (N = 36, from 20 families) for identification of novel and/or known pathogenic mutations. The clinical diagnosis was identified as either definite FH or possible FH using family history, individual lipid values, and history of tendon xanthomata in the patient or first degree family member, as well as information of segregation of high lipids in the family tree (1-3 generations). The number of individuals with definite clinical diagnosis were 22 (61%) from 11 families, and 14 individuals (39%) had possible clinical diagnosis from 11 families. The IHAFH study was approved by the National Bioethics Committee of Iceland (VSN-16-072-S2), and all participants gave written informed consent.

The cohort in the first phase of the Risk Evaluation For INfarct Estimates (REFINE) Reykjavik (18) was a random sample of 6940 men and women born in 1935-1985, living in Reykjavik area, between 20 and 74 years old with mean age of 49.6 years (SD 12 years). Age and sex specific cut-off values for the 95^th^ percentile in cholesterol were modelled using quantile regression. Models were run separately for the sexes, adjusted for age and restricted to individuals not on statins. The obtained predicted values were used as cut-offs to identify individuals in the REFINE cohort that were above the 95^th^ percentile in cholesterol stratified by age in groups of 5 years each (20-24, 25-29, 30-34, etc). Table 1 reports baseline characteristics of the REFINE Reykjavik study subjects. The REFINE Reykjavik study was approved by The National Bioethics Committee of Iceland (VSN-05-112-S1), and all participants gave written informed consent. Samples from both Icelandic cohorts were obtained from whole blood and the DNA isolated using a standard salting out method (20). The extracted DNA was mixed with a TE buffer in a tilt-rocking mixer for 24 hours and then stored at 4°C for a period of 4 weeks to ensure the DNA had completely dissolved into the TE buffer.

**Table 1:** Baseline characteristics of the study population. Baseline characteristics of the REFINE Reykjavik study cohort: Numbers are mean(SD) for continuous-, N(%) for categorical- and median[IQR] for skewed variables. Abbreviations: SBP, systolic blood pressure; DBP, diastolic blood pressure; TOT-C, total cholesterol; LDL-C, LDL cholesterol; TG, triglyceride; FG, fasting blood glucose; CHD, coronary heart disease; plaque, moderate or more plaque in carotids; N/A, not applicable.

| Characteristic | Variable | ≥ 95th percentile<br>TOT-C | <95th percentile<br>TOT-C | P-value* | Total<br>All subjects<br>in REFINE |
| --- | --- | --- | --- | --- | --- |
| **Demographics** |  |  |  |  |  |
|  | Numbers | 315 | 6625 |  | 6940 |
|  | Number females | 167 (53.0%) | 3363 (50.8%) | 0.435 | 3530 (50.9%) |
|  | Age (years) | 48.6 (11.7) | 49.7 (12.0) | 0.115 | 49.6 (12.0) |
| **Anthropometry** |  |  |  |  |  |
|  | BMI (kg/m<sup>2</sup>) | 28.0 (4.5) | 27.5 (5.0) | 0.042 | 27.5 (4.9) |
| **Physiological** |  |  |  |  |  |
|  | SBP (mmHg) | 124.3 (17.8) | 122.3 (16.6) | 0.039 | 122.4 (16.7) |
|  | DBP (mmHg) | 71.8 (9.7) | 71.1 (10.4) | 0.275 | 71.1 (10.3) |
|  | TOT-C (mmol/L) | 7.4 (0.8) | 5.1 (0.9) | <0.001 | 5.2 (1.0) |
|  | LDL-C (mmol/L) | 5.1 (0.8) | 3.1 (0.8) | <0.001 | 3.2 (0.9) |
|  | TG (mmol/L) | 1.7 [1.1,2.3] | 1.0 [0.7,1.4] | <0.001 | 1.0 [0.7,1.4] |
|  | FG (mmol/L) | 5.6 (1.3) | 5.5 (1.0) | 0.014 | 5.5 (1.0) |
| **Medication** |  |  |  |  |  |
|  | Antihypertension | 64 (20.3%) | 1601 (24.2%) | 0.112 | 1665 (24.1%) |
|  | Lipid lowering | 4 (1.3%) | 643 (9.7%) | <0.001 | 647 (9.3%) |
| **Lifestyle** |  |  |  |  |  |
|  | Smoker | 92 (29.5%) | 1367 (20.9%) | <0.001 | 1459 (21.3%) |
| **Metabolic** |  |  |  |  |  |
|  | T2D | 14 (4.4%) | 308 (4.6%) | 0.866 | 322 (4.6%) |
|  | Obese (BMI>30) | 86 (27.3%) | 1680 (25.4%) | 0.447 | 1766 (25.5%) |
| **Cardiovascular** |  |  |  |  |  |
|  | CHD prevalent | 5 (1.7%) | 252 (4.0%) | 0.047 | 257 (3.9%) |
|  | CHD incident | 21 (7.0%) | 357 (5.7%) | 0.345 | 378 (5.7%) |
|  | Plaque | 30 (9.5%) | 697 (10.5%) | 0.566 | 727 (10.5%) |

\*Comparing individuals >95<sup>th</sup> cholesterol percentile to the rest of the cohort. Obtained from two-sided T-test for continuous-, $\chi^2$ test for categorical- and quantile regression for skewed variable

For performance evaluation of the NGS panel we used samples of genomic DNA with previously identified variants in the genes interrogated by the sequencing panel that were obtained as purified DNA from the Human Genetic Cell Repository (National Institute of General Medical Sciences) at the Coriell Institute for Medical Research (Camden, NJ); the Indiana Biobank (Indianapolis, IN); the Euro Biobank (Milan, Italy); extracted from saliva samples collected by DNAsimple (Philadelphia, PA), or extracted from saliva samples through collaborations with the Hypertrophic Cardiomyopathy Association (Denville, NJ) and Sudden Arrhythmia Death Syndromes (Salt Lake City, UT) foundations.

All samples submitted to the Phosphorus Diagnostics laboratory as saliva were manually extracted using the Qiagen QIAamp Mini Kits following the manufacturer’s instructions. DNA was quantified using Quant-iT Picogreen dsDNA Assay Kit (Life Sciences) and a Varioskan LUX (Thermo Scientific). Supplementary Table S1 describes all validation samples used in this study, with biorepository numbers, as well as their previous characterization results and rationale for inclusion. All samples presented here are de-identified (HHS 45 CFR part 46.101(b)(4)). IRB approval to handle de-identified samples was obtained through ASPIRE (Santee, CA), protocol IRB-R-003.

### NGS panel description

A targeted next generation sequencing panel consisting of 179 genes related to cardiovascular disorders and sudden cardiac death, including cardiomyopathies, arrhythmias, aortopathies, pulmonary hypertension and lipidemias, was designed using a hybridization capture platform (Roche, Indianapolis, IN). Genes relevant to disease phenotype were included based on relationships described in Online Mendelian Inheritance in Man, Human Phenotype Ontology, GeneReviews, and primary literature. The panel included all coding exons, splice sites, promoter regions, 5’-Untranslated Regions (UTRs), and 3’-UTRs for each of the genes. Clinically relevant noncoding (intronic) regions that contained previously described pathogenic variants, as reported in ClinVar NIH database, were also included. The total size of the panel was 1,042,714 bp. Supplementary Table S2 shows the NGS panel gene content. For the present study, we focused on genes previously implicated in FH and ARH including the *LDLR*, *APOB*, *PCSK9* and *LDLRAP1* genes.

### Next Generation DNA Sequencing

All laboratory procedures were performed in a Clinical Laboratory Improvement Amendments (CLIA) laboratory (Phosphorus Diagnostics, New Jersey, NJ). DNA samples were prepared for sequencing using HyperPlus Library Preparation Kit (Roche, Indianapolis, IN) and sequenced on a NextSeq500 (Illumina, San Diego, CA), following the manufacturer’s instructions. A detailed sequencing protocol is provided in Supplementary Methods section.

### Variant classification

All bioinformatics algorithms were implemented within the Elements^™^ platform (Phosphorus, New York, NY). FASTQ files were produced from each sequencing run and processed using the germline calling pipeline (version 2.03.01.30066) in DRAGEN (Edico Genome, San Diego, CA). Variants identified by NGS were confirmed by an orthogonal method (microarrays or Sanger sequencing). After confirmation, each variant was classified as pathogenic, likely pathogenic, variant of unknown significance (VUS), likely benign or benign, following the American College of Medical Genetics (ACMG) guidelines (21).

### Orthogonal confirmation

A custom Phosphorus Affymetrix axiom array (22) was used to confirm all the reportable variants including SNVs, insertions/deletions <15 bp (indels) and CNVs. Microarrays were processed following the manufacturer’s instructions. To confirm variants not included in the microarray, Sanger sequencing was used. Orthogonal analysis was also performed for cases when there was discrepancy between expected (previously reported results) and obtained NGS results. If both NGS and confirmatory results agreed, the results were counted as concordant. Detailed protocols are described in the Supplementary Methods section.

## Results

### Analytical and clinical validation of the NGS gene panel

We designed a 179-gene panel including *LDLR*, *PCSK9*, *APOB* and *LDLRAP1*, as well as genes described in the literature that can harbor genetic defects involved in cardiac arrhythmias, cardiomyopathies, aortopathies and pulmonary hypertension. Gene content is described in Supplementary Table S2. In order to validate panel performance and determine analytical sensitivity, specificity and accuracy, we used 24 samples from the 1000 Genomes (1000G) project (23), for which location of SNVs and indels is known (Supplementary Table S1). NGS sequencing results from these validation samples was compared to known 1000G variants (Table 2). Microarray analysis and/or Sanger sequencing was performed to assess the discrepancies of SNVs/indels between results produced by the NGS panel and previously reported 1000G data. Analytical sensitivity of the test was >99% for SNVs and >94% for indels, and specificity was >99% for SNVs and >99% for indels. Final accuracy for SNVs and indels was 99.99% and 99.58%, respectively.

**Table 2:**
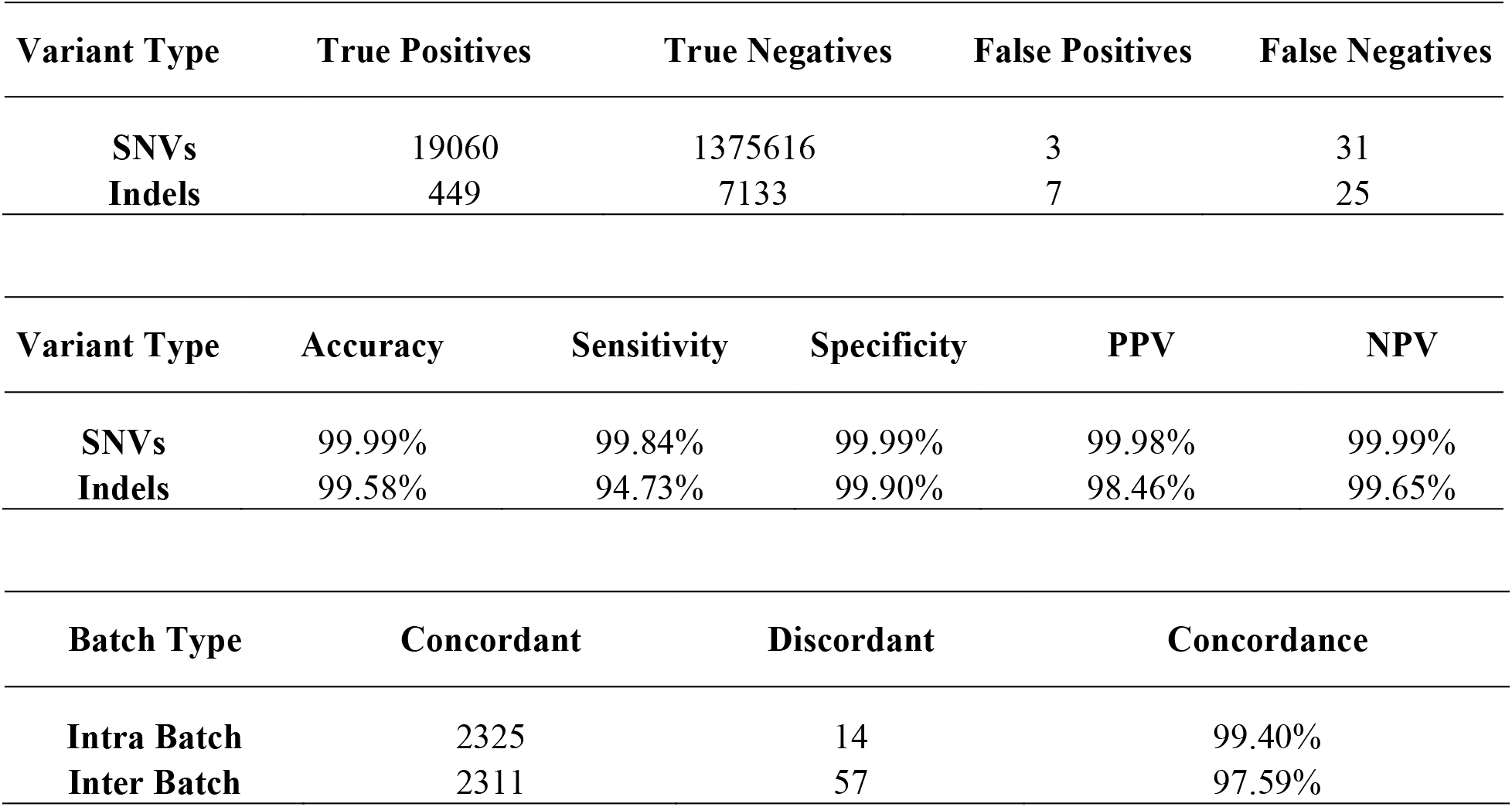
Performance characteristics of NGS panel. PPV, Positive Predictive Value. NPV, Negative Predictive Value.

| **Variant Type** | **True Positives** | **True Negatives** | **False Positives** | **False Negatives** |
| --- | --- | --- | --- | --- |
| **SNVs** | 19060 | 1375616 | 3 | 31 |
| **Indels** | 449 | 7133 | 7 | 25 |

| **Variant Type** | **Accuracy** | **Sensitivity** | **Specificity** | **PPV** | **NPV** |
| --- | --- | --- | --- | --- | --- |
| **SNVs** | 99.99% | 99.84% | 99.99% | 99.98% | 99.99% |
| **Indels** | 99.58% | 94.73% | 99.90% | 98.46% | 99.65% |

| **Batch Type** | **Concordant** | **Discordant** | **Concordance** |
| --- | --- | --- | --- |
| **Intra Batch** | 2325 | 14 | 99.40% |
| **Inter Batch** | 2311 | 57 | 97.59% |

Next, we determined clinical sensitivity. DNA samples harboring 61 known SNV/indels in the 179-gene panel were processed (N = 53; Supplementary Table S3). Known variants were confirmed in all the cases. Thus, clinical sensitivity for SNVs/indels by our assay was 100%. Furthermore, we identified three new pathogenic variants not reported previously in three different genes including, for instance, a premature termination mutation in *ALMS1* (subjects MQ39AT3, RTHC0315 and RTHC0418 Supplementary Table S3).

To determine clinical sensitivity in the detection of CNVs, we used samples with known CNV variants (N = 36; Supplementary Table S4). The set of previously known variants for these samples is limited, so specificity could not be assessed. The analysis correctly detected 32/36 CNVs, so the clinical sensitivity of the assay for CNVs is >88%. Importantly, the four CNVs missing are all due to the same 2502-nt deletion at NEB (NG_009382.2), spanning all of exon 55 plus flanking intronic sequences. This exon is only 104 bp long, a fragment that is of insufficient size to accurately perform CNV detection (Methods). An improved panel is currently being manufactured with added padding sequences in introns surrounding small exons to increase resolution in the detection of cases like this.

### Prospective testing by targeted panel screening in FH families

In order to determine diagnostic yield of the NGS panel in clinical samples, we screened for FH-causing genetic variants in two Icelandic cohorts: 1) 62 individuals from 23 FH families, of which 12 served as positive controls from three families with definite diagnosis of FH and with known culprit mutations, 14 individuals from 11 FH families that were included as negative controls as regards to clinical diagnosis, and 36 individuals from 20 families with definite or possible diagnosis of FH and unknown pathogenic mutations were sequenced using the NGS panel. Three mutations in the *LDLR* gene have previously been identified in the Icelandic population and were confirmed among the 12 positive controls (Supplementary Table S5): a 2kb deletion in the *LDLR* region of exons 9 to 10 (24), a SNV in exon 11 (NM_000527.4(LDLR):c.1618G>A (p.Ala540Thr)) and a SNV in intron/exon 4 (NM_000527.4(LDLR):c.694+2T>C) (19). Furthermore, two subjects from the Icelandic FH families and flagged as negative controls (subjects 93966177 and 94193565 in Supplementary Table S5) were found to carry pathogenic FH variants (Supplementary Figure 1) that co-segregated with FH among respective family members. One (subject 93966177) carried the splice donor variant c.694+2T>C in *LDLR* while the other (subject 94193565) carried a missense variant NM_000527.4(LDLR):c.1727A>G (p.Tyr576Cys), a novel FH pathogenic mutation in the Icelandic population (Supplementary Table S5). Upon closer inspection, these subjects were diagnosed at young age with high but borderline LDL-C and had older siblings with definite diagnosis of FH: subject 93966177 was a 15 yrs old male at the time of diagnosis with total cholesterol of 6.41 mmol/L and 5.15 mmol/L LDL-C, while subject 94193565 was an 8-yr-old female with total cholesterol of 4.92 mmol/L and 3.06 mmol/L LDL-C.

Among the 36 individuals with definite or possible diagnosis of FH and with unknown pathogenic mutations, we identified 16 subjects (from 6 families) with pathogenic/likely pathogenic variants (Supplementary Table S5). This included six FH variants in *LDLR* that segregated in different families and that have not been identified before among Icelandic FH families: NM_000527.4(LDLR):c.2120A>T (p.Asp707Val), NM_000527.4(LDLR):c.1727A>G (p.Tyr576Cys), NM_000527.4(LDLR):c.919G>A (p.Asp307Asn), NM_000527.4(LDLR):c.693C>A (p.Cys231Ter), 5.5 kb *LDLR* deletion removing part of exon 15 plus exon 16 and NM_000527.4(LDLR):c.1124A>G (p.Tyr375Cys) (Supplementary table S5). Thus, the diagnostic yield was 16/36 (44.4%) for all with diagnosis of FH, and 15/22 (68.2%) with definite diagnosis of FH (Supplementary Table S5). Overall, nine FH culprit mutations, all in the *LDLR* gene, have now been identified in the Icelandic population.

### Population-based prospective testing

We identified three individuals with FH pathogenic mutations (Supplementary table S6), among the 315 subjects in the top 95^th^ percentile cut-point for total cholesterol distribution in the REFINE Reykjavik study. One of these mutations was previously known in the Icelandic population, the 2kb deletion in the *LDLR* region of exons 9 to 10, while two (NM_000527.4(LDLR):c.693C>A (p.Cys231Ter)) and NM_000527.4(LDLR):c.1727A>G (p.Tyr576Cys) were novel in Iceland as described above. Thus, the prevalence of FH among the 315 with high cholesterol in the REFINE Reykjavik study was 0.95%. When we compared data from the whole REFINE Reykjavik cohort (N = 6940) with the Lipid Research Clinic FH register, 12 individuals were diagnosed with FH mutation in the REFINE Reykjavik study. Given REFINE Reykjavik is a population based study cohort we estimated the population frequency of FH to be at least 0.17% in Iceland. Thus, the enrichment of FH prevalence among those above the 95^th^ percentile in cholesterol is 5.6 fold (Fisher exact test P-value = 0.014). Note that the three FH patients among the top 5^th^ percentile in cholesterol levels had not been administered statins, while the other 9 FH patients received lipid lowering statin treatment. Finally, many of the 315 subjects were identified with VUS variants, most with conflicting classifications in ClinVar (https://www.ncbi.nlm.nih.gov/clinvar/) and some with discrepancies as likely pathogenic/VUS (Supplementary Table S7).

## Discussion

Early diagnosis and genetic testing of FH suspects is critical for improved prognosis, and the odds ratio for CHD adjusted for LDL cholesterol levels was 11.6 (95% CI 4.4-30.2) in individuals with clinical signs of FH and a confirmed culprit mutation (25). The effect of statin treatment on FH patients has been shown to be highly effective in reducing the risk of CHD that is comparable to the effect of lipid lowering in age-matched individuals from the general population (8).

Diagnosis of FH is usually done clinically by using a validated set of criteria (2, 9, 26). However, in children and young adults the disease is usually asymptomatic. Early diagnosis and treatment with statins is essential to reduce the risk of coronary heart disease (8, 27), therefore effective screening strategies should be available. FH increases in prevalence with age, peaking between ages 60 and 69 and declining thereafter (5). This suggests errors in diagnosis and insufficient dyslipidemia screening in children and adolescents, which was apparent for the two Icelandic children of FH families diagnosed free of the disease and marked as negative controls. This would suggest that screening for FH pathogenic mutations in young individuals at risk will enhance diagnosis of FH thus enabling early treatment and improved prognosis of the patients. Several genetic screening programs have been performed in different populations (28–30), however, it is still debated whether the increased clinical utility of identifying affected subjects justifies the costs of general and broad population genetic testing.

The screening of first-degree relatives of patients already diagnosed with FH and with further screening of first-degree relatives of the newly diagnosed relatives, has been referred to as cascade screening (31). Cascade screening in FH is cost effective (32), and supported by U.S. Centers for Disease Control and Prevention Office of Public Health Genomics (6). In Iceland, a genealogical database has been available from 1965 supporting extended cascade screening in very large families (33). Here, all index patients had a known FH mutation (19), and we showed that additional 19% of FH individuals were identified by using this extended family screening method compared with the use of the conventional cascade screening approach (33). A genetic diagnosis of the index patients is of great importance when these extended families are identified, in order to enhance the likelihood that the common ancestor (often born in the 18^th^ century) had been affected by this particular FH mutation and the screening for FH in their descendants to be successful. The identification of six new pathogenic variants in the *LDLR* gene in the Icelandic population can facilitate an extended cascade screening for identifying segregating culprit mutations for FH. Out of the 36 subjects clinically diagnosed with definite or possible FH, we identified 16 with pathogenic/likely pathogenic mutations offering a 44.4% diagnostic yield, while the diagnostic yield was 68.2% for those with definite diagnosis of FH. Thus, genetic testing through NGS is an excellent diagnostic method in cascade screening for FH pathogenic mutations.

Khera et al. analyzed the genes encoding *LDLR*, *APOB* and *PCSK9* in 956 individuals with LDL-C ≥ 190 mg/dl from 5 population cohort studies (34). The prevalence of FH confirmed in these individuals was 1.7%, while the population prevalence of FH was 0.13% in these cohorts, resulting in a 13-fold enrichment of FH in the target sample. The inclusion criteria were stricter in Khera et al. than in the present study, as only individuals with very high LDL-C were selected and lipid lowering treatment was considered, whereas we selected individuals based on percentiles total cholesterol by age and gender. Therefore, our study included more of younger individuals with borderline levels of high cholesterol. In spite of these differences, we found that the prevalence of FH was 0.95% in the group of 315 with total cholesterol above the 95^th^ percentile cut-point compared to the FH population prevalence of 0.17%, thus providing a significant 5.6 fold enrichment in FH. Note that the three FH patients among the top 5^th^ percentile in cholesterol levels had not been administered statins, while the other 9 FH patients in REFINE received lipid lowering treatment. Therefore, it is likely that statin treatment masked the hypercholesterolemia phenotype thus excluding some FH patients for entering the top 5^th^ percentile group for total cholesterol. In the present study, we have demonstrated that genetic testing by NGS may be cost effective as a diagnostic method in a population-based screening for FH pathogenic mutations. However, the diagnostic yield is sensitive to the stratification of the target population.

A novel NGS-based targeted panel for the genetic diagnosis of FH was validated which also included a number of candidate genes involved in an array of other cardiovascular related diseases. Similar to other reported screening panels (16, 17), parameters like sensitivity, specificity and accuracy of the present panel proved to be excellent, showing that NGS is a technically mature platform that is useful for confident genetic screening and diagnosis of rare disease like FH. However, there are remaining caveats as regards variant classification and interpretation (35), including for instance in fields like hypertrophic cardiomyopathy and heart arrhythmias. This is of lesser concern for FH, due to the vast amount of data accumulated over the years on the functional impact of variants in genes involved in the LDL receptor pathway. Cholesterol levels in FH and non-FH subjects can overlap considerably and especially in adults, thus it would be an advantage if lipid disorder NGS panels that included both FH and non-FH genes were made available.

In summary, genetic diagnosis of FH by NGS is feasible and informative for cascade screening of family members, as well as providing a significant diagnostic yield for the general population screening. Also, the present study suggests diagnostic yield could be enhanced in population screening for FH by proper stratification of the target population. Advances in sequencing technologies are occurring at fast pace, which will help lower the cost and, at some point, facilitate identification of subjects at risk at young age, therefore allowing early pharmacological intervention and improvement prognosis of affected individuals.

## Acknowledgments

We thank the study subjects and family members who consented to participate in this research project. We thank Ed O’Neill for editorial comments on the manuscript.

## Abbreviations

ARH: Autosomal Recessive Hypercholesterolemia
CNV: Copy Number Variant
FH: Familial Hypercholesterolemia
LDLR: LDL-receptor gene
LDL-C: LDL cholesterol
NGS: Next Generation Sequencing
MLPA: Multiplex Ligation-dependent Probe Amplification
SNV: Single Nucleotide Variant
UTRs: Untranslated Regions
VUS: Variant of Unknown Significance

